# Surface Chemistry and Interfacial Biophysics of Meibum and Meibum—Benzalkonium Chloride Films: Implications for Tear Film Lipid Layer

**DOI:** 10.64898/2026.06.13.731656

**Authors:** Jianzhong Chen, Yiming Zhang, Thai Minh Han Nguyen, Vladimir V Tsukruk

## Abstract

The tear film lipid layer (TFLL) prevents excessive evaporation, thereby maintaining normal ocular surface function. One characteristic of a healthy TFLL is a gel-like structure that resists disruptive mechanical forces; however, this has not been clearly demonstrated. Herein, the compressibility of TFLL was investigated via surface chemistry experiments of meibomian gland secretions (meibum), the source of TFLL. Meibum was collected post-mortem from bovine eyelids, and purity was verified by mass spectrometry. Meibum films on a water subphase in a Langmuir-Blodgett trough were studied. The changes in surface pressure (*π*) versus area (*A*) during compression were recorded, and the corresponding compressive modulus (*C*_s_^-1^) versus *π* isotherms were determined. The meibum film was stable at a surface pressure as high as 32 mN/m and maintained a near-constant *C*_s_^-1^ of 24–32 mN/m across a π range of 8–30 mN/m, demonstrating gel-like behavior. Comparison of the maximum *C*_s_^-1^ with those of representative individual meibum lipids supports the presence of a monolayer predominantly composed of low-polarity lipids, including wax esters, cholesteryl esters, and diesters. The average molecular areas determined for the film and reported for representative lipid species suggest that the monolayer is predominantly composed of saturated, ultralong-chain, low-polarity lipids and that the TFLL includes a dense monolayer and a loose overlying sublayer. This structural model is further supported by the lower *C*_s_^-1^ (10 mN/m) and larger average molecular area (68 Å^2^) observed for films containing a mixture of meibum and benzalkonium chloride (BAK), a preservative with surfactant properties.

## 1. Introduction

A thin tear film covers and lubricates the ocular surface to maintain normal visual function. The tear film is structurally organized into an outermost lipid layer, a middle aqueous layer, and an innermost mucin layer, with the latter two often collectively referred to as the muco-aqueous layer. The superficial tear film lipid layer (TFLL) serves as a critical barrier that prevents excessive evaporation of the underlying aqueous phase (Willcox et al., 2017).

One characteristic of a healthy TFLL is proposed to be a gel-like, incompressible structure that resists disruptive mechanical forces (King-Smith et al., 2013). This characteristic can be studied by the compressive modulus (*C*_s_^-1^) versus *π* isotherms of the TFLL, where *C*_s_^-1^ data points are determined from surface pressure (*π*) versus surface area (*A*) isotherms (Davies and Rideal, 1963; Oliveira et al., 2022). As the source of TFLL and readily available in greater amounts, meibum samples have been used in these surface chemistry and interfacial biophysics studies. Meibum samples studied include those from humans (Brown and Dervichian, 1969; Butovich et al., 2010; Leiske et al., 2010; Georgiev et al., 2011; Mudgil and Millar, 2011; Leiske et al., 2012a; Leiske et al., 2012b), cows (Brown and Dervichian, 1969; Petrov et al., 2007; Mudgil and Millar, 2008; Georgiev et al., 2010; Leiske et al., 2010), and other mammals (Leiske et al., 2010; Yoshida et al., 2019). However, this proposed gel-like TFLL characteristic has not been clearly demonstrated.

This gel-like, incompressible structure of the TFLL can be further tested using *C*_s_^-1^ versus *π* isotherms of a film of a mixture of meibum and a surfactant such as benzalkonium chloride (BAK). BAK is used as a preservative at a concentration of 0.005-0.02% in 70% of commercially available eye drops, with a major side effect of dry eye, typically attributed to cytotoxicity (Goldstein et al., 2022; Kahook et al., 2024). As a surfactant, BAK should be able to insert into the TFLL and even displace lipid molecules at the air-water interface. However, the effects of BAK on the surface pressure of lipid layers tested by surface chemistry are ambiguous (Georgiev et al., 2011; Riedlová et al., 2023). One reason could be that BAK was prepared in an aqueous solution in the form of micelles, and the preformed lipid layer reduces the penetration of BAK into the air-water interface, slowing the displacement of lipid molecules by BAK to the point that it is undetected within minutes or hours in a typical Langmuir experiment. Therefore, studying the surface chemistry and compressibility of the film formed by direct mixing of BAK and meibum is likely to shed light on TFLL structure and function.

In this study, we collected post-mortem bovine meibum, prepared meibum lipid solutions in chloroform, both alone and mixed with 10% BAK, verified their purity by mass spectrometry and liquid chromatography, and then performed surface pressure measurements during compression in a Langmuir-Blodgett trough. The *π*-*A* and *C*_s_^-1^-*π* isotherms were determined to assess the surface chemistry and interfacial biophysics of meibum film and the effect of BAK.

## 2. Methods and materials

### 2.1. Chemicals and reagents

Chloroform (CHROMASOLV Plus, ≥99.9%, amylenes as stabilizer), methanol (CHROMASOLV, LC-MS grade, >99.9%), ammonium hydroxide (≥25% in water, for LC-MS, Fluka^TM^), and water (CHROMASOLV, LC-MS) were purchased from Fisher Scientific (Waltham, MA, USA). Benzalkonium chloride (≥95.0%) was purchased from Sigma (St. Louis, MO, USA).

### 2.2. Bovine meibum collection and stock solution preparation

One pair of upper and lower eyelids from one eye of each of 3 cows, obtained from the slaughterhouse at FPL Food LLC (Augusta, GA), was used in this study. Bovine meibum was collected by expressing the eyelids (**Fig. 1**) using a method similar to that previously reported (Nicolaides et al., 1981). Briefly, individual eyelids were warmed to 37°C in a water bath, cleaned with ultrapure water, squeezed with pliers to express meibum, and then collected into a microcapillary glass tube or scooped into a glass vial with a spatula. A total of 4.9 mg of meibum collected from one cow, combining secretions from both upper and lower eyelids of a single eye, was dissolved in chloroform to prepare a stock solution of 5 mg/mL. This large amount of meibum allowed accurate weight measurements and preparation of the stock solution sufficient for both mass spectrometry analysis and Langmuir trough experiments. The stock solution was stored at −80°C until ready for experiments.

**Fig. 1.**
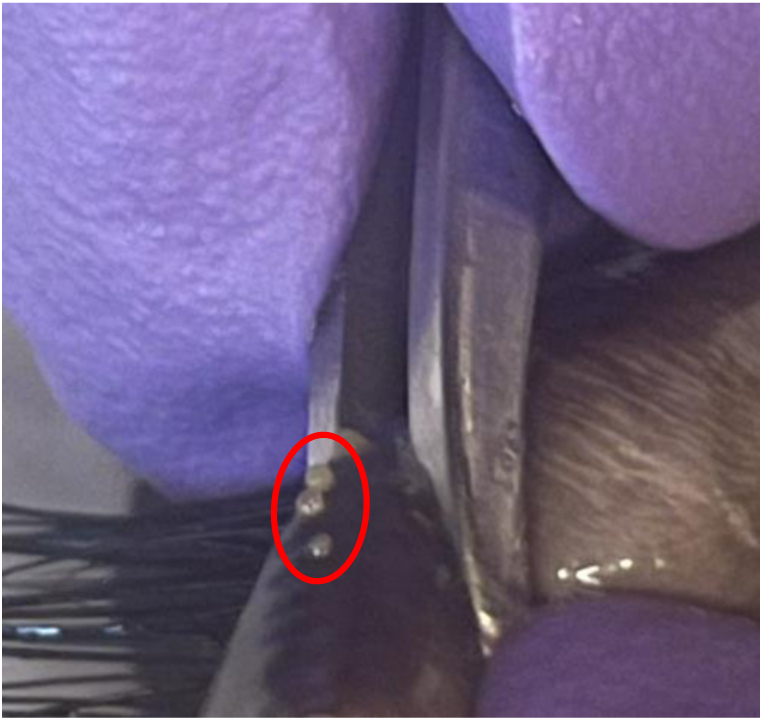
Expression of meibum from a bovine eyelid. Three droplets of meibum at the gland orifices are circled in red.

### 2.3. Lipidomics analysis of bovine meibum

The lipid composition of bovine meibum was analyzed by shotgun lipidomics (Han and Gross, 2005; Züllig et al., 2020) using a method (Chen et al., 2025) adapted from our previous method (Chen and Nichols, 2018) to an Orbitrap Fusion Tribrid mass spectrometer equipped with an electrospray ionization source (Thermo Fisher Scientific, Waltham, MA, USA). Bovine meibum working solutions for mass spectrometry analysis were diluted from the stock solution to final concentrations of 9 µg/mL and 90 µg/mL containing 0.1% and 1% ammonium hydroxide in chloroform-methanol solvent mixtures of 1:99 (v/v) and 10:90 (v/v), respectively. The meibum lipid solutions were infused into the mass spectrometer at a flow rate of 5 µL/min. Mass spectrometer parameters include: capillary temperature 175°C; vaporization temperature 25°C; and spray voltages of 3500 V and 3000 V in positive-and negative-ion modes, respectively. The ammonium hydroxide additive was previously found to work effectively in both positive-and negative-ion modes (Chen and Nichols, 2018; Chen and Panthi, 2019). For tandem mass spectrometry, high-energy collisional dissociation (HCD) at a normalized collision energy of 5%, 20%, 30%, 40%, or stepped collision energy (15%, 30%, and 45%) was applied via sequential precursor-ion fragmentation, as previously reported by our group (Chen et al., 2025). Experiments under each condition were performed in duplicate.

### 2.4. Langmuir-Blodgett experiments

The isotherm experiments were performed in a KSV 2000 Langmuir-Blodgett Minitrough (KSV Instruments Ltd, Helsinki, Finland) with two movable barriers, using a Wilhelmy plate to measure surface pressure under different compression regimes, following a procedure described in detail earlier (Flouda et al., 2022). The Teflon trough and barriers were cleaned with chloroform and ultrapure water (18.2 MΩ·cm; Synergy UV-R, EMD Millipore). Experiments were conducted at room temperature (20°C). Ultrapure water was used as the subphase. The working solutions of bovine meibum for surface chemistry analysis were prepared by diluting the stock solution to 1 mg/mL in chloroform, with one solution additionally containing 0.1 mg/mL BAK. A 200 µL of each solution was overlaid onto the water subphase dropwise and left undisturbed for 30 minutes to allow chloroform to evaporate. The lipid layers were compressed by moving the two barriers toward each other at 5 mm/min for each until they reached the minimum barrier separation distance or collapsed. After expansion, second compressions were performed. The surface pressure (*π*) changes upon compression of the lipid film with area (*A*) were recorded. The *C*_s_^-1^ at each data point was determined from the equation *C*_s_^-1^ =-*A*(d*π*/d*A*) (Davies and Rideal, 1963; Georgiev et al., 2011; Guaus and Torrent-Burgués, 2018), where *A* is the surface area at a specific *π*.

## 3. Results

### 3.1. Mass spectrometry analysis of lipid molecular species in bovine meibum

Mass spectrometry analysis of bovine meibum samples in positive and negative ion modes showed a similar pattern across the two concentrations. The main difference is that compared to the high-concentration sample spectra, the lower-concentration sample spectra contained negligible noncovalently bound dimer peaks, which were formed in the gas phase under the mass spectrometry experimental conditions (**Figs. 2a & 2b**). The bovine meibum were predominantly composed of wax esters (WEs), cholesteryl esters (CEs), and two types of diesters (DEs), along with a small amount of other lipids, including triacylglycerols (TGs) and cholesterol, detected in positive ion mode (**Fig. 2a**). A small amount free fatty acids (FFAs) and O-acyl-ω-hydroxy fatty acids (OAHFAs) were detected in negative ion mode (**Fig. 2b**). The identities of these lipids were confirmed by HCD tandem mass spectrometry (**Fig. 3**). The predominant WEs in bovine meibum include those with the fatty alcohol/fatty acid moiety combination being 25:0/15:0, 27:0/15:0, 25:0/18:1, and 27:0/18:1. The predominant CEs include CE 25:0 and CE 27:0. The most abundant triacylglycerol is 52:2. Few impurity peaks were observed (**Figs. 2 & 3**).

**Fig. 2.**
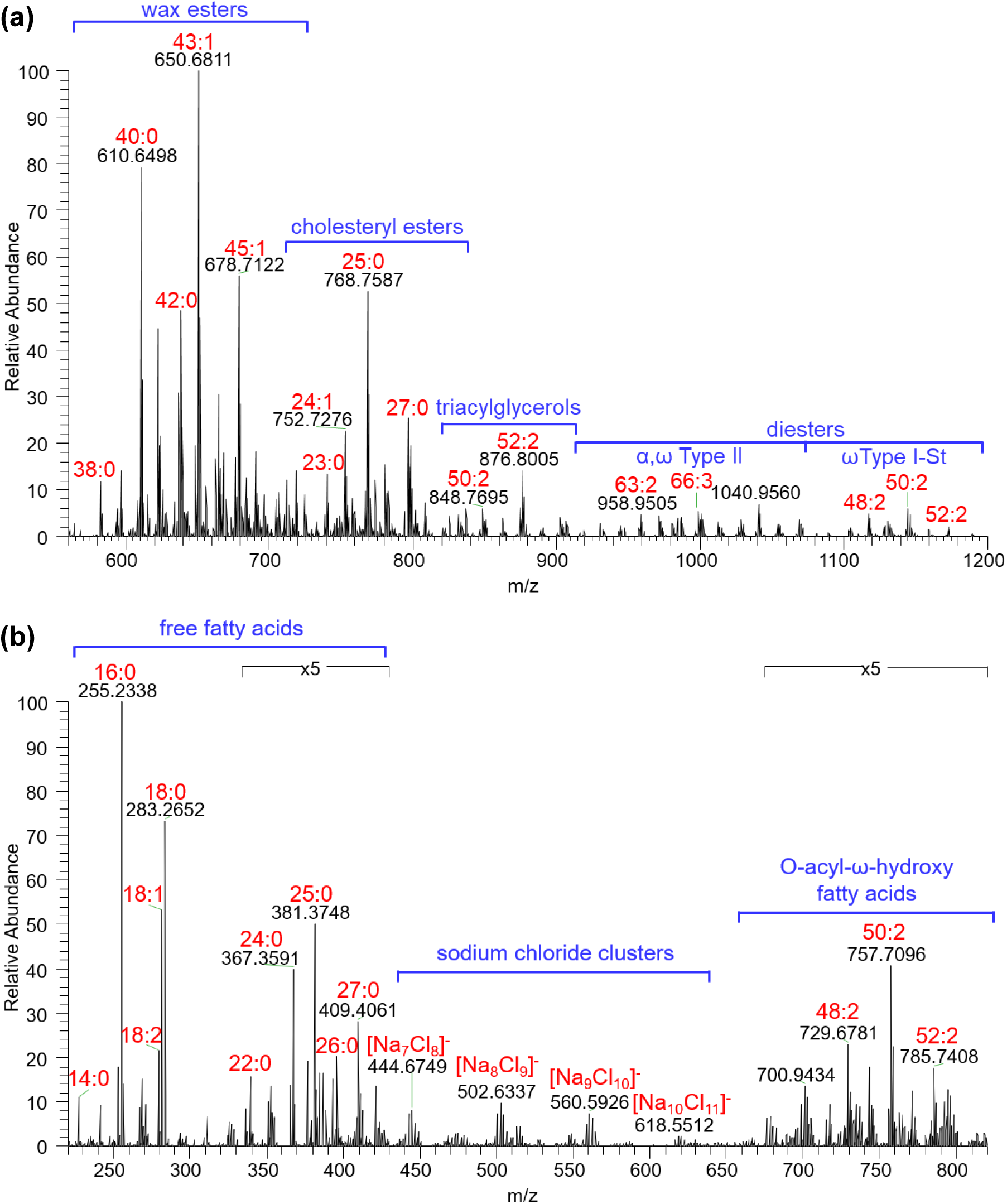
Mass spectra of bovine meibum. **a)** positive ion mode; and b) negative ion mode. The cholesterol peak (*m*/*z* 369.3521), not shown in a), was detected at 8.5% of the highest intensity peak (*m*/*z* 650.6811). The two numbers separated by a colon that label peaks for wax esters, α,ω Type II diester species, free fatty acids, and O-acyl-ω-hydroxy fatty acids are, respectively, the total number of carbon atoms and the number of double bonds. For cholesteryl esters and ω Type I-St diesters, the total number of carbon atoms and the number of double bonds of the fatty acyl substituent are shown. For triacylglycerols, the total number of carbon atoms and the summed number of double bonds of the three fatty acyl chains are shown.

**Fig. 3.**
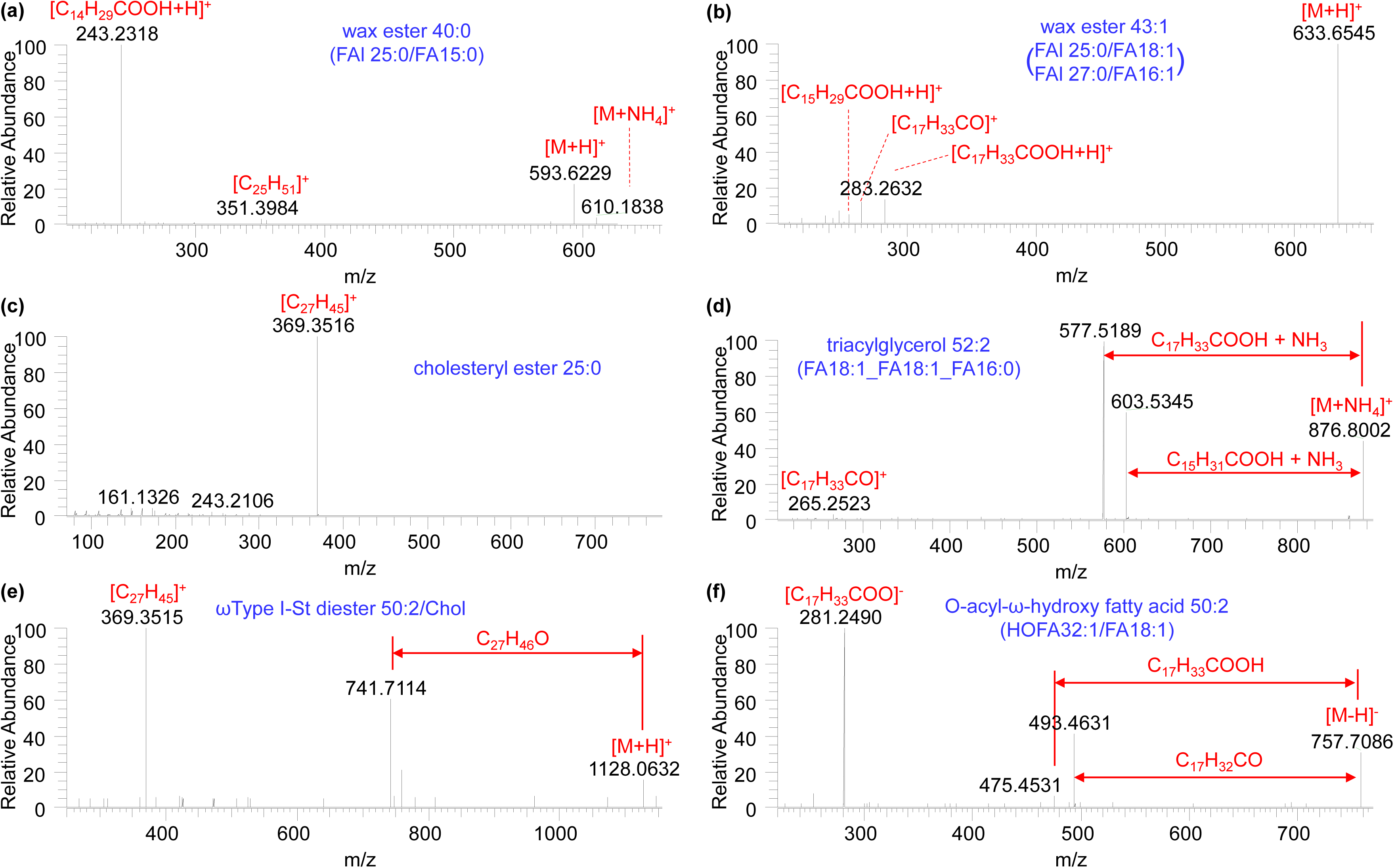
Collision-induced MS/MS spectra of representative lipid molecular species in bovine meibum. **a)** Wax ester 40:0; b) wax ester 43:1; c) cholesteryl ester 25:0; d) triacylglycerol 52:2; e) ωType I-St diester 50:2/Chol; and f) O-acyl-ω-hydroxy fatty acid 50:2. Each molecular species may comprise more than one isomer, and the abundant component(s) for each panel are shown in parentheses. Abbreviations: Chol, cholesterol; FAl, fatty alcohol; FA, fatty acid; HOFA, hydroxy fatty acid.

### 3.2. π ‒ A and C_s_^-1^ ‒ π isotherms of meibum-only films

The *π* ‒ *A* isotherms for bovine meibum films demonstrate that the lipid layer was stable up to ∼32 mN/m (**Figs. 4a**). Consistent with the gel-like structure proposed for the TFLL (King-Smith et al., 2013), the *C*_s_^-1^ ‒ *A* and *C*_s_^-1^ ‒ *π* isotherms for the meibum sample (**Figs. 4b & 4c**) included a wide region of constant *C*_s_^-1^ (24-32 mN/m), which became narrower (19-25 mN/m) during the second compression (**Figs. Suppl. 1b & 1c**). However, the incompressible structure proposed for the TFLL (King-Smith et al., 2013) appears inaccurate; in contrast, the low *C*_s_^-1^ indicates the TFLL is highly compressible (Davies and Rideal, 1963). The plateau region of *π* and reduced *C*_s_^-1^ at the end of the first compression (**Fig. 4a**) is likely due to the film approaching collapse at certain spots. Some fluctuation in the *C*_s_^-1^ ‒ *A* and *C*_s_^-1^ ‒ *π* isotherms (**Figs. 4b, 4c, Suppl. 1b & 1c**) was due to the complexity of the meibum lipid composition and reorganization of the film.

**Fig. 4.**
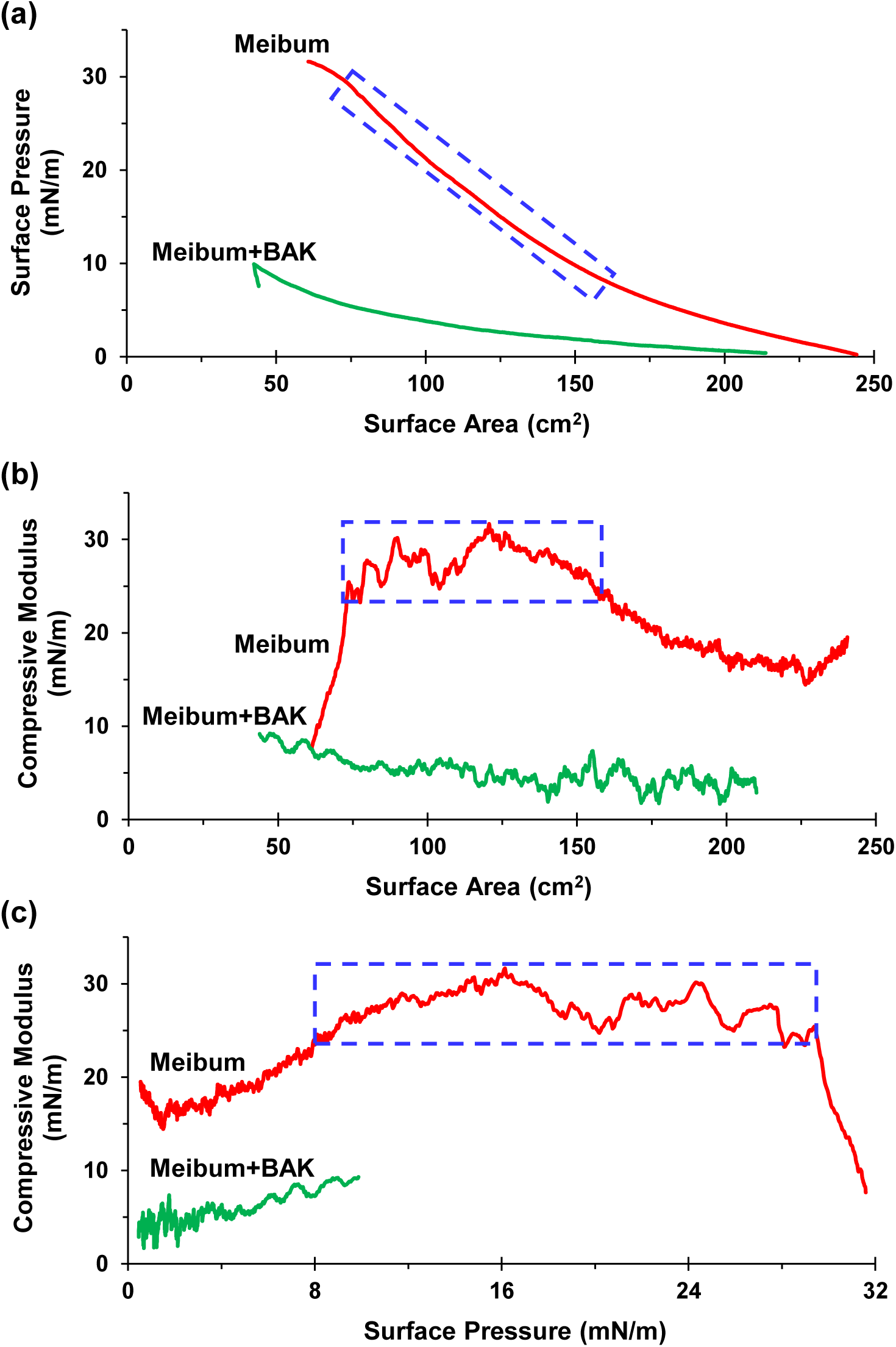
Isotherms for meibum and meibum-BAK films on water. **a)** surface pressure (*π*) vs. surface area ( *A*); b) compressive modulus (*C_s_^-1^*) vs. surface area (*A*), and c) compressive modulus (*C_s_^-1^*) vs. surface pressure (*π*). The constant-compressive-modulus regions are indicated by dashed rectangles.

### 3.3. π ‒ A and C_s_^-1^ ‒ π isotherms of meibum–BAK mixture films

The meibum-BAK mixture films collapsed at a surface pressure as low as ∼10 mN/m ( **Fig. 4a & Suppl. 1a**). The *C*_s_^-1^ values also dropped sharply, with the maximum slightly below 10 mN/m and no constant *C*_s_^-1^ regions were detected in the *C*_s_^-1^ ‒ *π* isotherms (**Fig. 4c** & **Supplementary Fig. 1c**). The *π* ‒ *A* isotherms of the second compressions appeared largely reproducible with those of the first compressions (**Suppl. Fig. 1a** & **Fig. 4a**). In contrast, the *C*_s_^-1^ ‒ *A* and *C*_s_^-1^ ‒ *π* isotherms for the second compressions showed a memory effect of the first compressions: the *C*_s_^-1^ values were initially close to those at the end of the first compression; they then first gradually decreased, then increased, until reaching a plateau (**Suppl. Fig. 1b & 1c**).

### 3.4. Average areas per molecule for meibum and meibum-BAK mixture films

The average molecular weight of lipid molecules in meibum was estimated to be 700 ± 12 Da. It is determined from the molar concentrations and corresponding molecular weights of WE and CE molecular species (Chen et al., 2013). WEs and CEs account for 88% ± 5% of total meibum lipids (Chen et al., 2013). Thus, the estimated molecular weight is close to the true value.

The average molecular weight of BAK was determined to be 344.5 Da using the percentages of its individual components and their corresponding molecular weights. The batch of BAK we received included 70.9% benzyldimethyldodecylammonium chloride (C_12_-BAK) and 28.1% benzyldimethyltetradecylammonium chloride (C_14_-BAK).

The *π ‒ A* isotherms, with *A* in units of area per molecule, were then plotted based on the putative molecular composition of the monolayers at the air-water interface (**Fig. 5**). The average areas per molecule were calculated assuming that the monolayers contain 1) all meibum lipid molecules, 2) all saturated, ultralong-chain (SULoC)-containing WE and CE molecules, and 3) all BAK molecules, respectively. Please see *4.4.* and *4.5.* for detailed discussions.

**Fig. 5.**
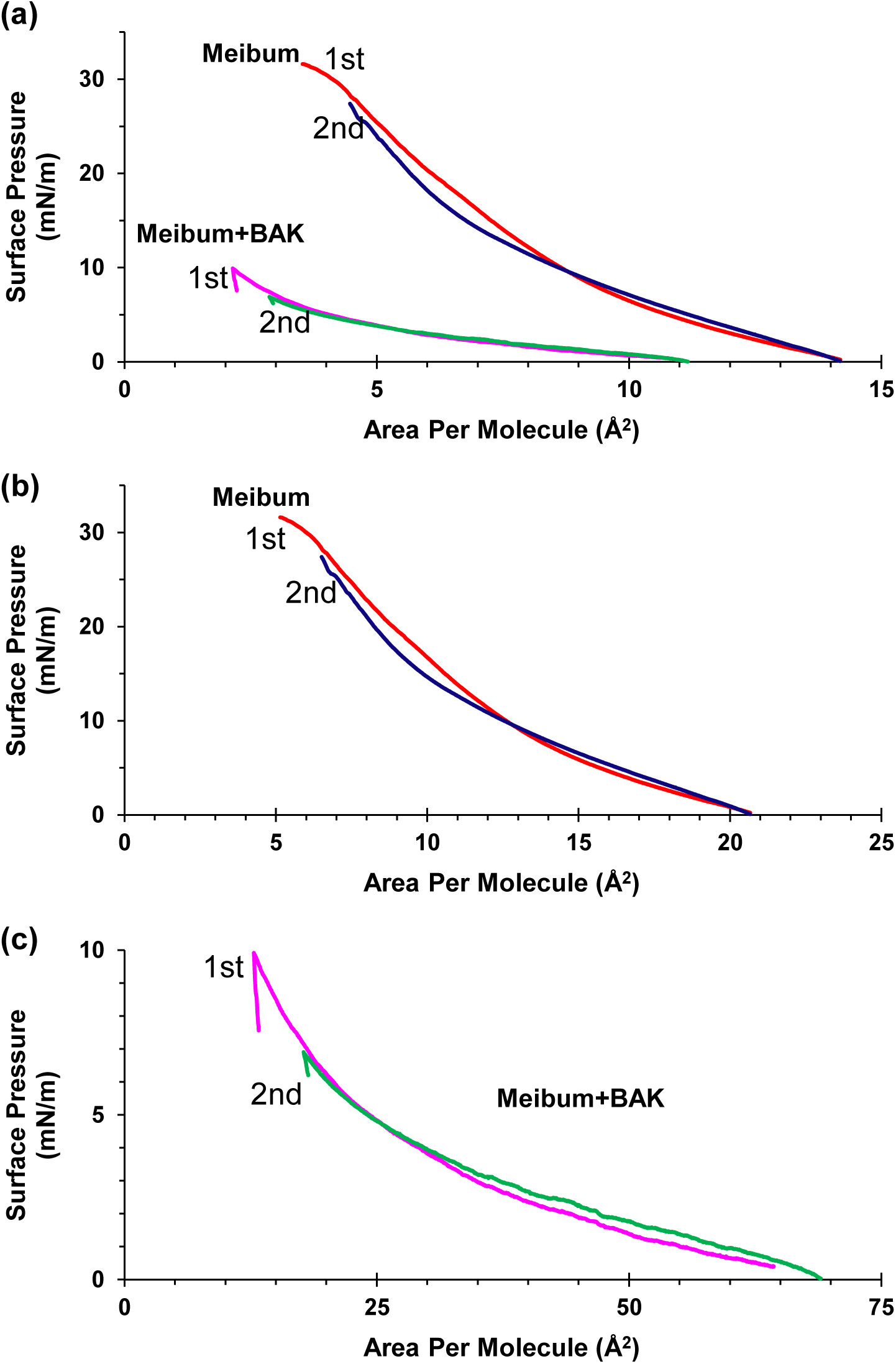
Surface pressure (*π*) vs. area per molecule (*A*) isotherms for meibum-only and meibum-BAK films on water. Shown are plots assuming: a) all molecules formed a monolayer; b) only saturated ultralong-chain (SULoC)-containing wax esters and cholesteryl esters formed a monolayer; and c) only BAK molecules formed a monolayer.

## 4. Discussion

### 4.1. Lipid molecular species in bovine meibum

To our knowledge, this is the first mass spectrometry analysis of intact lipid molecules in bovine meibum. The intact lipid composition of bovine meibum (**Fig. 2**) is similar to that reported for human meibum (Chen et al., 2010; Chen and Nichols, 2018; Chen et al., 2019). The predominant lipid components in meibum from both species are WEs, CEs, DEs, along with TGs, cholesterol, FFAs, and OAHFAs, while phospholipids and sphingomyelins are negligible. Nevertheless, some differences are also apparent. The ultralong chains in WEs and CEs are approximately one carbon atom longer in bovine meibum than in human meibum (Chen et al., 2013, 2016), consistent with previous reports (Nicolaides et al., 1981; Kolattukudy et al., 1985). This may be related to the need for greater evaporation resistance due to the larger exposed corneal surface area of a bigger eye (Howland et al., 2004). This is consistent with a previous report that evaporation resistance increases exponentially with the chain length of saturated fatty alcohols (La Mer and Healy, 1965). Another difference between human and bovine meibum is that the fatty acyl chain 15:0 in WEs is more abundant in bovine than in human (**Fig. 3a**). This is consistent with the previous report that the FA15:0 moiety (summed from normal-, *iso*-, and *anteiso*-chains) in WEs accounts for 40.80% of the total fatty acyl moiety for bovine meibum versus 1.48% for human meibum (Nicolaides et al., 1981). In contrast, the most abundant fatty acyl chain in human WEs is FA18:1, accounting for 56.57% of the total, whereas in bovine meibum it is only 21.01% (Nicolaides et al., 1981). The reason for the different combinations of fatty acyl and fatty alcohol moieties in WEs is likely the need to maintain a subtle balance of intermolecular interactions, enabling both high evaporation resistance and high re-spreadability, two characteristics that seem incompatible (King-Smith et al., 2013).

Our results show both differences and similarities with previous reports (Baron and Blough, 1976; Nicolaides et al., 1981). The major components of meibum identified across these studies are the same, including WEs, CEs, DEs, TGs, cholesterol, and FFAs. The unidentified “post-TG region” lipids in the references (Nicolaides et al., 1981) are likely OAHFAs (**Fig. 2b**), which were not identified until the 2010s (Butovich et al., 2009; Chen et al., 2010). The moieties in WEs and CEs are also similar in these studies, predominantly of saturated ultralong-chain (Cn = 25-27) fatty alcohol (FAl) moieties in WE and fatty acid (FA) moieties in CE (**Figs. 2 & 3**) (Baron and Blough, 1976; Nicolaides et al., 1981). For instance, the summed amount for fatty acyl and fatty alcohol chains 25:0 and 27:0 (normal-, *iso*-, and *anteiso*-chains combined) was >50% in CEs and >75% in WEs of bovine meibum, respectively (Nicolaides et al., 1981). In contrast, polar lipids, with a high percentage (13.26%) reported in (Nicolaides et al., 1981) or 1.7% in (Baron and Blough, 1976), were not detected in our results (**Figs. 2 & 3**). Likewise, high-percentage polar lipids (16.04%) reported for human meibum (Nicolaides et al., 1981) were not detected by modern mass spectrometry (Butovich et al., 2009; Chen et al., 2010). The quantitative information from our results also appears to differ from that of these earlier studies: the abundance of WEs and CEs is higher, while TGs are lower. These earlier reports identified and quantified meibum lipids using methods including fractionation, saponification, derivatization, and gas chromatography coupled to a flame ionization detector or an electron ionization mass spectrometer for separation and detection; thus, only chain moieties were determined. In contrast, our methods allowed us to identify the chain combination in intact lipids, which gained insight into intermolecular interactions. A more detailed investigation of the lipid composition of bovine meibum is ongoing in our laboratory.

### 4.2. π ‒ A isotherms of meibum films compared to literature reports

Previous studies reported a wide range of maximum values of *π* (from 10 to 49 mN/m) for *π* ‒ A isotherms of meibum and tears (**Table 1**). Nevertheless, the value of 32 mN/m in our study is in excellent agreement with a narrower range of 33 to 37 mN/m reported in 6/15 of previous bovine and human meibum studies (**Table 1**). Importantly, our value is consistent with the value of 28 ± 3 mN/m for normal human tears, calculated from the surface tension of 43.6 ± 2.7 mN/m (n=65, ranging from 38.4 to 52.1 mN/m) (Tiffany et al., 1989) and water surface tension of 72 mN/m at 25°C (Vargaftik et al., 1983). The smooth curves in our study (**Figs. 4a & Suppl. 1a**) also agree well with reports on bovine meibum (Petrov et al., 2007), human meibum (Georgiev et al., 2011), and human tears (Guaus and Torrent-Burgués, 2018).

**Table 1.**
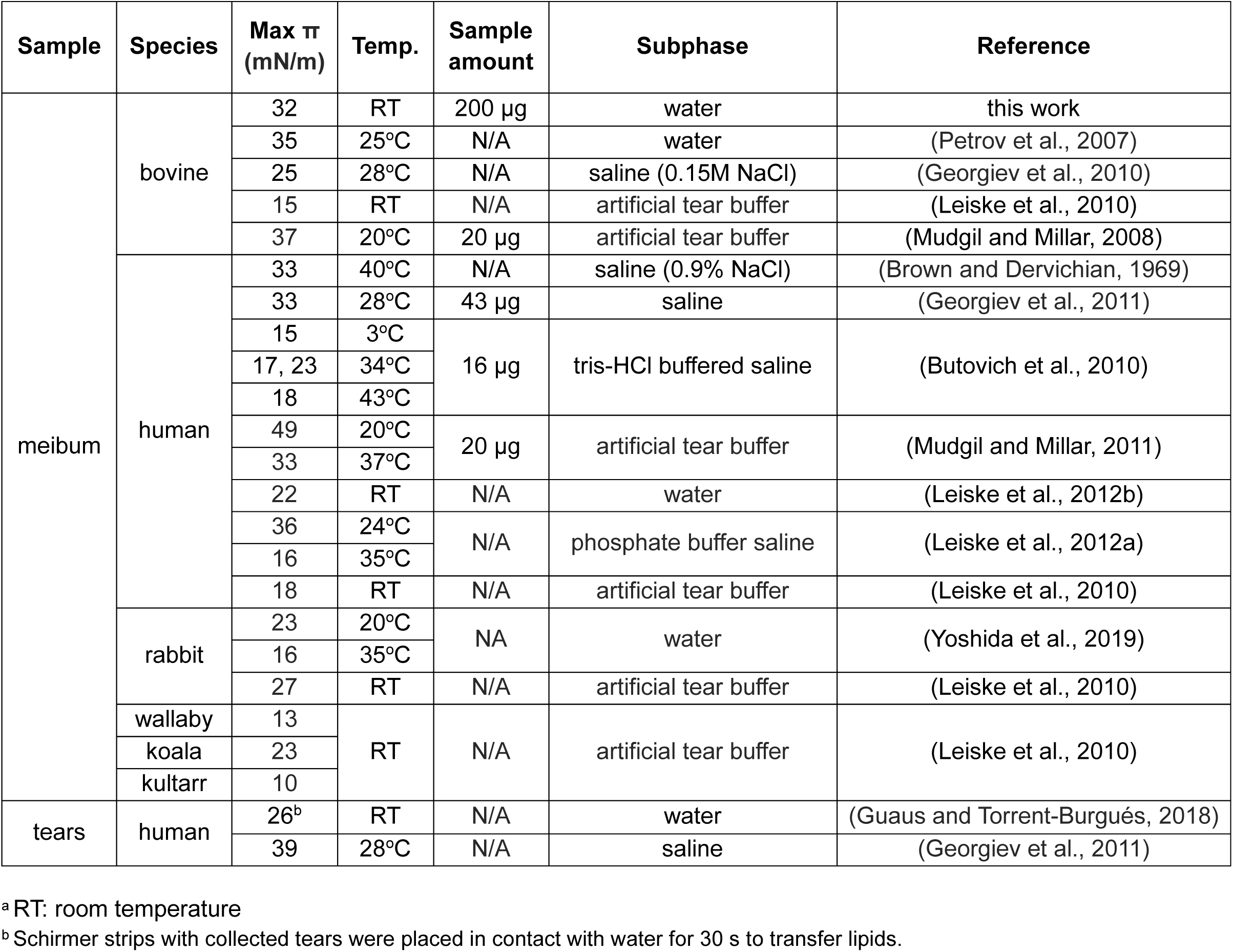
List of maximum surface pressure (π) of meibum and tears lipid layers^a^.

A quasi-plateau region at the end of compression in our experiments (**Fig. 4a**) was not reported in most previous studies, including two reports with nearly the same maximum *π* value (Petrov et al., 2007; Georgiev et al., 2011). One explanation is that these two studies compressed the monolayer at higher speeds, possibly resulting in metastability (Smith et al., 2003). The compression speed was 11 mm/min in (Georgiev et al., 2011); it was not mentioned in (Petrov et al., 2007) and was assumed to be 0.8 Å^2^/(molecule·min), the same as in a paper by the same group (Petrov et al., 2000). In contrast, our compression speed was 5 mm/min, corresponding to 0.4 Å^2^/(molecule·min), only about half of the two studies. Also, we loaded a greater amount of lipids into the Langmuir trough, ranging from 5-fold to more than 10-fold that of the other studies (**Table 1**), which allowed us to monitor the isotherms over a broad range and accurately determine the area per molecule. As mentioned in the introduction, a lower loading amount decreases the maximum *π* reached (Butovich et al., 2010).

Other variables in experiments for the *π‒A* isotherms include contamination, subphase, and temperature. Common contaminants include phospholipids from dead corneal cells during sample collection (Kunnen et al., 2016; Chen et al., 2019), plasticizers during sample processing, and palmitic acid and stearic acid from unknown sources (Ende and Spiteller, 1982; Keller et al., 2008; Chen, 2021). Compared to the maximum *π* for the individual meibum lipid class (**Table 2**), phospholipids and FFAs tend to increase the maximum *π* for the meibum film while plasticizers and TGs tend to decrease the maximum *π*. In contrast, the subphase composition did not appear to significantly affect the isotherms (**Table 1**). This is likely because the TFLL or meibum film includes a critical monolayer at the air-aqueous layer interface comprised predominantly neutral lipids that are not influenced by pH (Chen et al., 1997; Chen et al., 2013; Chen and Panthi, 2019) (see *4.4.* for details). The effect of temperature, on the other hand, is ambiguous. One study showed that *π* increased with temperature (Butovich et al., 2010); three others showed the opposite trend (Mudgil and Millar, 2011; Leiske et al., 2012a; Yoshida et al., 2019), however, at different levels (**Table 1**).

**Table 2.**
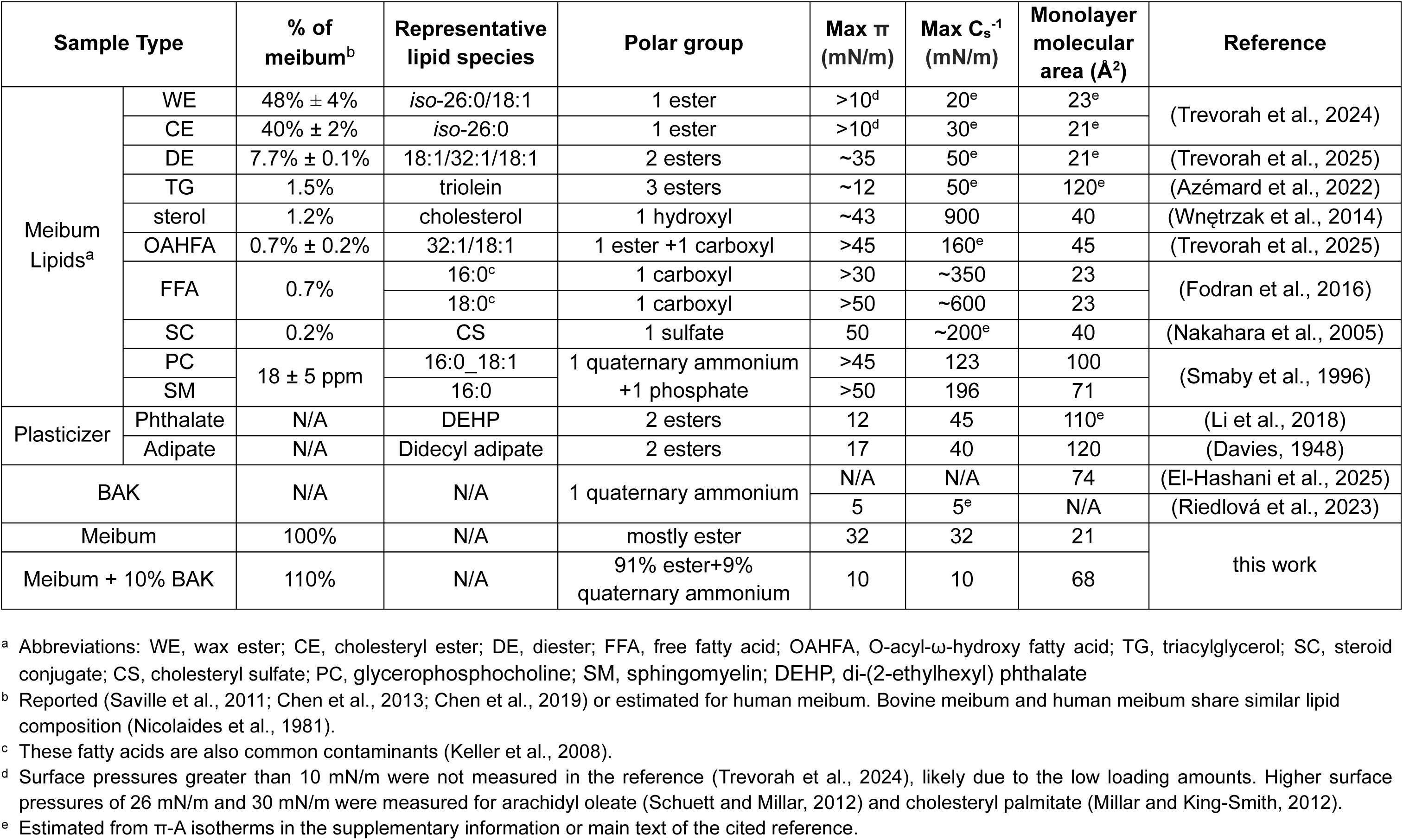
Surface properties determined in this work versus those of major meibum lipid classes and plasticizers.

### 4.3. C_s_^-1^ ‒ π isotherms of meibum films compared to literature reports

A constant *C*_s_^-1^ region (24-32 mN/m) across *π* range of 8-30 mN/min was observed in this study (**Fig. 4c**). Similar observations of constant *C*_s_^-1^ regions were reported previously, including 28 mN/m across *π* range of 25-33 mN/m for human meibum (Georgiev et al., 2011), and 29-31 mN/m across *π* range of 7-12 mN/m for human tears (Guaus and Torrent-Burgués, 2018) (**Table 3**). These studies reported similar *C*_s_^-1^ values; however, the regions were much narrower. One reason for the wider constant *C*_s_^-1^ region in our study could be the lower barrier-moving speed (5 mm/min, corresponding to 7.5 cm²/min) compared to 11 mm/min in (Georgiev et al., 2011) and 30 cm²/min in (Guaus and Torrent-Burgués, 2018). This lower speed allowed lipid molecules to have sufficient time to organize into highly ordered, uniform phases. Compared with the human tear study (Guaus and Torrent-Burgués, 2018), a greater amount of bovine meibum lipids in our study resulted in a higher *π*. Interestingly, the constant region in the study of human tears appears smoother than in the two meibum studies, likely because high-polarity lipids in the tear sample were scavenged by tear lipocalin (Glasgow et al., 1995) and possibly by other lipid-binding proteins such as lacritin (Georgiev et al., 2021), and thus less interference to the monolayer by these lipids (see *4.4.* for more detailed discussion).

**Table 3.** List of constant compressive modulus (C_s_^-1^) and the corresponding surface pressure (π) ranges of meibum and tears lipid layers.

| Sample | Species | Temperature | Constant $C_s^{-1}$ region | | Reference |
| --- | --- | --- | --- | --- | --- |
| | | | $C_s^{-1}$ (mN/m) | $\pi$ (mN/m) | |
| meibum | bovine | RT | 24-32 | 8-30 | this work |
|  | human | 28°C | ~28<sup>a</sup> | 25-33 | (Georgiev et al., 2011) |
| tears | human<sup>b</sup> | RT | 29-31 | 8-9.5 | (Guaus and Torrent-Burgués, 2018) |

<sup>a</sup> Estimated from a $C_s^{-1}$ - $\pi$ isotherm on a logarithmic scale for $C_s^{-1}$ .
<sup>b</sup> collected on Schimmer strips and placed in contact with water for 30 s.

### 4.4. Implications for the TFLL structure by comparing C_s_^-1^ values of meibum, tears, and individual meibum/tear lipids

Comparing the biophysical properties of the lipid layers of meibum, tears, and individual meibum/tear lipid components may shed light on the lipids at the air-aqueous layer interface. All the high-polarity meibum/tear lipid classes, including glycerophosphocholine, sphingomyelin, cholesterol, O-acyl-ω-hydroxy fatty acids, and free fatty acids, display high maximum *C*_s_^-1^ values (100-900 mN/m). Therefore, it is unlikely that these high-polarity lipids would, by themselves, form a monolayer in the TFLL at the water-air interface, as previously suggested (McCulley and Shine, 1997; Rosenfeld et al., 2013). In contrast, major nonpolar lipids, including WEs, CEs, and DEs, showed values of maximum *π* (>10 to 50 mN/m) and *C*_s_^-1^ (20-50 mN/m) comparable to those of meibum (25-33 mN/m) and *C*_s_^-1^ (24-32 mN/m) (**Table 2**). Although these lipids are often called nonpolar lipids, they contain one or two polar ester groups (Sadgrove and Jones, 2019) that could interact with water molecules. Thus, “low-polarity lipids” should be the accurate term. Importantly, these low-polarity lipids account for more than 95% of total meibum lipids (Chen et al., 2013). Therefore, it is more likely that these low polarity lipids interact coordinately with water molecules at the water-air interface, supporting our hypothesis that the TFLL comprises a dense monolayer (Chen and Panthi, 2019).

Literature reports further suggest that of the low-polarity lipids, saturated ultralong-chain species comprise the monolayer. Of note, human meibum contains exactly the same saturated ultralong chains 24:0, 25:0, and 26:0 (fatty alcohol moieties in WEs and fatty acid moieties in CEs) in the top 3 abundant WEs (Lam et al., 2013; Chen et al., 2016) and CEs (Nicolaides et al., 1981; Chen et al., 2010; Butovich, 2010; Chen et al., 2013; Brown et al., 2013). A similar pattern is also observed in bovine (see discussion in **3.1**) and tree shrew meibum (Chen and Panthi, 2019). The nonpolar carbon chains of the same or different classes can interact with each other via dipole-dipole interactions (Flory and Vrij, 1963). While the dipole-dipole interaction is weak between common carbon chains, it increases with the length of linear carbon chains. The linear parts of these saturated ultralong chains (or equivalent) are >1.5 times longer compared to typical chains in other lipids or other tissues (18 carbon atoms with a kink at the 9^th^ carbon atom) (Chen and Panthi, 2019), allowing formation of a stable monolayer between the carbon chains of these classes (Kotz, 2019). These results strongly suggest that saturated, ultralong-chain (SULoC)-containing WEs and CEs interact to form a monolayer (**Fig. 6**). In contrast, the unsaturated, ultralong-chain-(UULoC-) counterparts contain double bonds that introduce a kink in the chain, thereby shortening the linear part of the chain. The shorter chains lead to weaker intermolecular forces, as suggested by melting point studies (Knothe and Dunn, 2009). These lower forces would be unable to hold these lipids together at the air-water interface in a hairpin shape, as the SULoC low-polarity lipids do. As a result, these lipids are likely squeezed out above the monolayer (La Mer and Aylmore, 1962), where they adopt a relaxed, extended conformation (**Fig. 6**). Our ongoing molecular dynamics studies will test the hypothesis that SULoC low-polarity lipids form a stable monolayer.

**Fig. 6.**
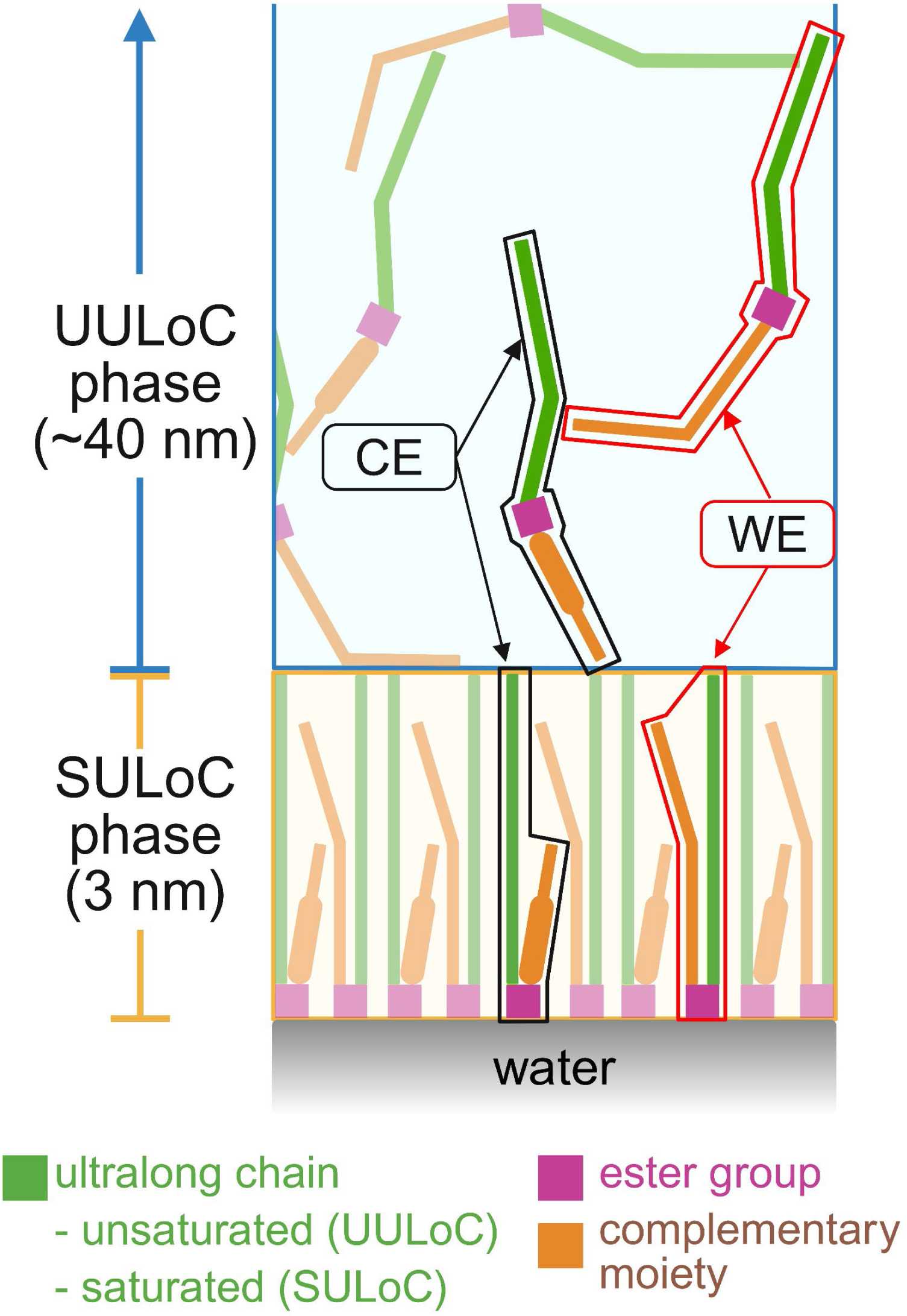
Proposed two-sublayer structure and composition for meibum lipid film overlaid on water. Saturated ultralong-chain (SULoC)-containing wax esters and cholesteryl esters form a monolayer at the air-water interface, while the unsaturated ultralong-chain (UULoC)-containing counterparts have shorter linear chains (due to a kink introduced by a double bond), resulting in weaker intermolecular interactions and thus being squeezed above the monolayer. For clarity, other lower-abundance lipids are not shown.

Triacylglycerols display a *C*_s_^-1^ value (45 mN/m) slightly higher than that of meibum and tears, and in theory, they could participate in the monolayer in terms of *C*_s_^-1^. However, they lack ultralong chains and possess 3 crowded ester groups, which may make it difficult for them to interact with WE and CE molecules (Chen and Panthi, 2019). On the other hand, with 3 ester groups, their polarity is indeed higher than that of WEs, CEs, and DEs. Therefore, we classify them as high-polarity lipids.

This two-sublayer TFLL structure — one dense monolayer and one loose outer sublayer — was previously observed in the rat TFLL via an *in vivo* cryofixation with freeze-substitution (VC-FS) electron microscopy (Chen et al., 1997). Nevertheless, their conclusion that the rat tear film lacked a free aqueous layer is inaccurate. This conclusion was based on the network-like structure beneath the lipid layer and the assumption that the lipid layer completely prevents evaporation. However, the network structure was simply the remaining salts and proteins after evaporation of water — the lipid layer only slows, rather than prevents, evaporation.

The two-sublayer structure is further supported by the determined areas per molecule. When assuming all meibum lipid molecules formed a monolayer, the average molecular area was 14.2 Å^2^ (**Fig. 5a**). Compared to the molecular areas (21-23 Å^2^) of the individual low-polarity lipids (**Table 2**), the number of lipid layers correspond to approximately one and a half if these lipids were well-packed. If assuming SULoC WEs and CEs (likely along with DEs containing chains equivalent to SULoC) form a monolayer, the average molecular area is 20.9 Å^2^ (**Fig. 5b**), which is consistent with 21-23 Å^2^ for WEs, CEs, and DEs (**Table 2**). The calculation was based on that SULoC WEs and CEs account for ∼78% each of the two lipid classes (**Table 4**), while WEs and CEs account for 88% of total meibum lipids (Chen et al., 2013); taken together, these SULoC lipids sum to 68% of total meibum lipids. Our preliminary data suggest SULoC-equivalent-containing DEs account for ∼82% of total DEs, while DEs account for 7.7% of total meibum lipids (Chen et al., 2013). When these lipids are included in the monolayer, the corresponding average molecular area is 19.1 Å^2^, still very close to molecular areas of WEs, CEs, and DEs in a monolayer. The slightly lower molecular area is also consistent with our molecular dynamics simulation results (collaborating with Vance Jaeger, PhD, manuscript in preparation) that these lipids are slightly more tightly packed than WEs alone. High-polarity meibum lipids, including cholesterol, OAHFA, FFA, cholesteryl sulfate, PC, and SM, which could have been scavenged by lipocalin in tears, would remain in the monolayer of the meibum film in this study. However, these lipids account for <5% of total meibum lipids and therefore will not significantly affect the calculation. UULoC(and equivalent)-containing WEs and CEs, as well as TGs, accounting for 21% of meibum lipids, would then form a ∼10-fold thicker loose outer sublayer (∼40 nm) (Willcox et al., 2017). The corresponding density would be approximately 3% of that of the monolayer, consistent with the VC-FS electron microscopy study of the rat TFLL (Chen et al., 1997). This two-sublayer TFLL structure hypothesis is also consistent with the high-resolution color microscopy studies showing that the human TFLL was non-homogeneous, with the thinnest part being only 1-2 molecules thick (King-Smith et al., 2011; Millar and King-Smith, 2012).

**Table 4.**
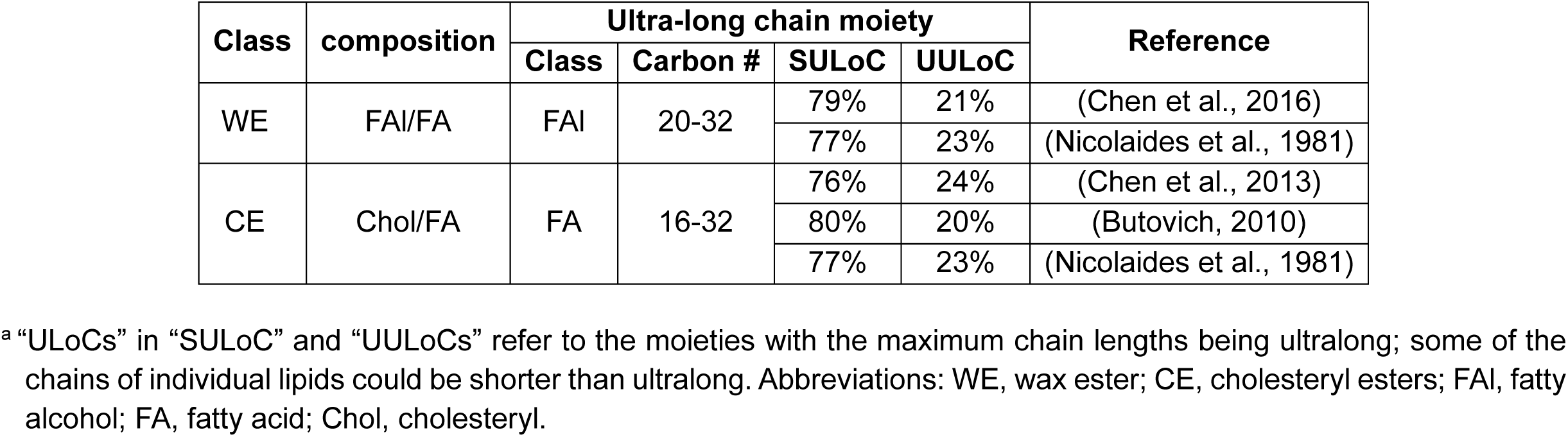
Relative amount of saturated ultra-long-chain (SULoC) and unsaturated ultra-long-chain (UULoC) moieties in human meibum wax esters and cholesteryl esters^a^.

### 4.5. π ‒ A and C_s_^-1^ ‒ π isotherms of meibum-BAK mixture films

Compared to the meibum films, both maximum *π* and *C*_s_^-1^ values of the meibum-BAK mixture films were drastically reduced from 32 mN/m to < 10 mN/m (**Figs. 4 & Suppl. Fig. 1**). A *C*_s_^-1^ of 12.5 – 50 mN/m, 100 – 250 mN/m, and 1000 – 2000 mN/m corresponds to liquid expanded, liquid condensed, and solid condensed states, respectively (Davies and Rideal, 1963). These surface chemistry experiments demonstrate that the meibum-only lipid layers are in a gel-like, liquid-expanded state with constant compressibility over a wide range of surface pressure. In contrast, the meibum-BAK mixture films’ *C*_s_^-1^ values are approximately equal to those of the surface pressure (**Fig 4c & Suppl. Fig. 1c**), characteristic of an ideal gas (Davies and Rideal, 1963).

The lower *π* and *C*_s_^-1^ for the meibum-BAK mixture films suggest BAK forms a monolayer at the air-aqueous interface. More than 95% of meibum lipids are WEs, CEs, and DEs (Chen et al., 2013), which contain only one or two low-polarity ester groups. As discussed above, in the absence of a large amount of high-polarity lipids, these saturated-ultra-long-chain-containing species of these lipids likely form a stable lipid layer (Chen and Panthi, 2019). However, in the mixture of meibum and 10% BAK, BAK contains a head group (quaternary ammonium) that is much more polar (Sadgrove and Jones, 2019) than the ester groups of the predominant lipids in meibum. The high polarity and large amount of BAK allowed BAK molecules to interact strongly with water molecules to form a monolayer while squeezing the meibum lipids above the aqueous-lipid interface (La Mer and Aylmore, 1962). Indeed, when assuming that all BAK molecules form a monolayer, the determined area per molecule (68 Å^2^) (**Fig. 5c**) is consistent with the reported BAK molecular area (74 Å^2^) (El-Hashani et al., 2025). Meanwhile, BAK contains one short benzyl group, one long carbon chain (predominantly 12, 14, or 16 carbon atoms) (Liu et al., 2017), and two methyl groups; all these groups are hydrophobic and covalently bound to the same nitrogen atom. This structure tends to orient the cation horizontally in an extended conformation, leaving no leeway for alternative orientations, such as those of an ester group, and thus leading to lower *π* and greater compressibility (lower *C*_s_^-1^). A similarly low maximum π (< 5 mN/m) was previously reported for an aqueous BAK solution at 0.36 µM with an 8-carbon chain in a phosphate-buffered saline subphase (Riedlová et al., 2023). The collapse of the meibum-BAK mixture film under compression at low π (< 10 mN/m) is likely due to BAK cations forming micelles at a locally high concentration (Kato et al., 2013).

### 4.6. Implications for BAK-induced dry eye mechanism

As a preservative in eye drops, BAK is dissolved in water in a micellar form (0.005-0.02%) (Goldstein et al., 2022; Kahook et al., 2024) because of its low critical micelle concentration, i.e., 0.0025-0.15 mM, corresponding to 0.00001-0.005% (Kato et al., 2013). To mimic this situation, previous reports studied the surface chemistry of meibum or tear lipid layers by injecting a BAK aqueous solution into the subphase (Georgiev et al., 2011). With a preformed lipid layer, these BAK cations were blocked from adsorbing at the water-air interface and were likely instead adsorbed onto the measuring Wilhelmy Plate beneath the lipid layer, thereby increasing the overall *π* to > 40 mN/m. A similar phenomenon of increased *π* in the presence of BAK solution was also reported for mixtures of representative lipids (Riedlová et al., 2023). However, given sufficient time, BAK could insert into the adsorbed meibum lipid layer through strong interactions with water molecules and replace part of the meibum lipids, or, under extreme conditions, all of them. Another major difference between clinical conditions and the surface chemistry experiments is that the tear film’s aqueous layer is only 3-4 µm while human eyes blink ∼20 times per minute, which leads to very good mixing of BAK with the aqueous layer. In contrast, for the surface chemistry experiments, 250 µL BAK solution was injected into 40 mL subphase (Georgiev et al., 2011), which may take quite some time to result in efficient mixing.

The preparation of the BAK-meibum mixture solution in chloroform in this study differs from the typical clinical condition, in which an aqueous BAK solution is instilled in the presence of a preformed TFLL. Nevertheless, the mechanism for BAK disrupting meibum film could be the same as that for the side effect of long-term use of BAK-containing ophthalmic eye drops, although at different levels. The observed reduced *π* and *C*_s_^-1^ (**Fig. 4**) may represent the worst-case equilibrium scenario for these clinical applications.

To investigate the typical dry eye side effect level of BAK in ophthalmic eye drops in patients, further surface chemistry and interfacial biophysics studies are warranted. These studies may include mixing different percentages of BAK with meibum, mxing individual BAK components of different chain lengths (Liu et al., 2017; Riedlová et al., 2023) with meibum, mixing BAK with subphase effectively first before depositing meibum solution, or long-term equilibration of aqueous BAK solution with a preformed TFLL. These studies will provide insight into this understudied mechanism for the adverse effects of BAK.

Meibum contains a small amount of high-polarity lipids, including fatty acids, cholesterol, and phospholipids (**Fig. 2**) (Chen et al., 2010; Saville et al., 2011; Chen et al., 2013). These polar lipids could have long-term effects similar to those proposed for BAK. However, except for phospholipids, the polarity of these lipids is much lower than that of BAK. Although phospholipids contain a high-polarity quaternary ammonium group that likely disrupts the low-polarity lipid layer, their concentrations in meibum and tears are very low, below 20 ppm (Saville et al., 2011). Moreover, almost all of these high-polarity lipids are known to bind lipocalin, a carrier protein (Glasgow et al., 1995), and thus can be scavenged from the air-water interface. In contrast, there is no such mechanism to remove BAK from the water-air interface. The permanent charge on BAK makes it interact strongly with water molecules noncovalently and is likely to remain at the water-air interface for a long time rather than be excreted with the TFLL, potentially causing a problem in the TFLL.

## 5. Conclusions

This study supports the hypothesis that the TFLL comprises a dense monolayer of low-polarity lipids at the air-water interface and a loose outer lipid sublayer. Comparing *C*_s_^-1^ of meibum films and individual meibum lipids reported in the literature (**Table 2**) suggests that only a monolayer predominantly composed of WEs, CEs, and DEs could form a monolayer with a *C*_s_^-1^ comparable to that of meibum or TFLL. Consistently, the average area per molecule of the monolayer composed of SULoC WEs, CEs, and DEs (accounting for ∼75% of total meibum lipids) was determined to be 19-21 Å^2^, matching the value (21-23 Å^2^) of these esters. While most of the remaining low-polarity meibum lipids, accounting for approximately 20% of the lipids in the monolayer, composes a loose outer lipid layer with a volume more than 10-fold that of the monolayer. The remaining high-polarity lipids, accounting for 5% of total meibum lipids, most likely scavenged into the bulk aqueous layer in the tear film under physiological conditions, would remain in the monolayer in the Langmuir-Blodgett experiments. Nevertheless, the small amount of these lipids would not significantly change *C*_s_^-1^ and the average area per molecule of the monolayer.

Our team’s collaborative work will further test this hypothetical TFLL structure using a variety of methods to analyze meibum, tears, and their fractions. These methods include: 1) lipid layer transfer from a liquid subphase to a solid substrate in a Langmuir-Blodgett (LB) trough (Oliveira et al., 2022; Buxton et al., 2025), 2) surface characterization of the transferred monolayer via spectroscopic ellipsometry (Politano and Versace, 2023; Bukharina et al., 2024) and high-resolution atomic force microscopy (Tsukruk and Singamaneni, 2012; Bukharina et al., 2024), and 3) all-atom molecular dynamics simulations (Khanom et al., 2026). Confirming this hypothetical structure will provide critical insights to guide future research into the mechanisms, diagnosis, and treatment of dry eye disease.

## Acknowledgements

The authors thank Dr. Pranita Patil from FPL Food LLC for providing bovine eyelids, Dr. Wenbo Zhi from Augusta University for technical assistance with the mass spectrometer, Dr. Shruti Sharma from Augusta University for providing benzalkonium chloride, and Dr. Christine Curcio from the University of Alabama at Birmingham for a critical reading of the manuscript.

## Abbreviations

BAK: Benzalkonium chloride
Chol: cholesterol
CE: cholesteryl ester
DE: diester
FA: fatty acid
FAl: fatty alcohol
HOFA: hydroxy fatty acid
TFLL: tear film lipid layer
WE: wax ester

---

**Suppl. Fig. 1. Isotherms for meibum and meibum-BAK films on water upon the second compression.** a) surface pressure (*π*) vs. surface area (*A*); b) compressive modulus (*C_s_^-1^*) vs. surface area (*A*), and c) compressive modulus (*C_s_^-1^*) vs. surface pressure (*π*). The constant-compressive-modulus regions are indicated by dashed rectangles.

## Notes

### Competing Interest Statement

The authors have declared no competing interest.

### Summary of Updates

This version of the manuscript has been revised to include the monolayer molecular areas of meibum, meibum-BAK films, and representative meibum lipids, and the proposed two-sublayer structure and composition for meibum lipid film overlaid on water.

### Funding

- NEI Center Core Grant for Vision Research: P30EY031631

## References

Azémard, C., Fauré, M.C., Stankic, S., Chenot, S., Ibrahim, H., Laporte, L., Fontaine, P., Goldmann, M., de Viguerie, L., 2022. Influence of unsaturations on the organization and air reactivity of triglyceride monolayers. Langmuir 38, 711–718.

Baron, C., Blough, H.A., 1976. Composition of the neutral lipids of bovine meibomian secretions. J. Lipid Res. 17, 373–376.

Brown, S.H., Kunnen, C.M., Duchoslav, E., Dolla, N.K., Kelso, M.J., Papas, E.B., Lazon de la Jara, P., Willcox, M.D., Blanksby, S.J., Mitchell, T.W., 2013. A comparison of patient matched meibum and tear lipidomes. Invest. Ophthalmol. Vis. Sci. 54, 7417–7424.

Brown, S.I., Dervichian, D.G., 1969. The oils of the meibomian glands: Physical and surface characteristics. Arch. Ophthalmol. 82, 537–540.

Bukharina, D., Southard, L., Dimitrov, B., Brackenridge, J.A., Kang, S., Min, P., Wang, Y., Nepal, D., McConney, M.E., Bunning, T.J., Kotov, N.A., Tsukruk, V.V., 2024. Left and right-handed light reflection and emission in ultrathin cellulose nanocrystals films with printed helicity. Advanced Functional Materials 34, 2404857.

Butovich, I.A., 2010. Fatty acid composition of cholesteryl esters of human meibomian gland secretions. Steroids 75, 726–733.

Butovich, I.A., Arciniega, J.C., Wojtowicz, J.C., 2010. Meibomian lipid films and the impact of temperature. Invest. Ophthalmol. Vis. Sci. 51, 5508–5518.

Butovich, I.A., Wojtowicz, J.C., Molai, M., 2009. Human tear film and meibum. Very long chain wax esters and (o-acyl)-omega-hydroxy fatty acids of meibum. J. Lipid Res. 50, 2471–2485.

Buxton, M.L., Brackenridge, J., Poliukhova, V., Nepal, D., Bunning, T.J., Tsukruk, V.V., 2025. Surface mapping of functionalized two-dimensional nanosheets: Graphene oxide and mxene materials. Langmuir 41, 11866–11881.

Chen, H.B., Yamabayashi, S., Ou, B., Tanaka, Y., Ohno, S., Tsukahara, S., 1997. Structure and composition of rat precorneal tear film. A study by an in vivo cryofixation. Invest. Ophthalmol. Vis. Sci. 38, 381–387.

Chen, J., 2021. Mass spectrometric analysis of meibomian gland lipids, in: Hsu, F.-F. (Ed.), Mass spectrometry-based lipidomics: Methods and protocols. Springer US, New York, NY, pp. 157–170.

Chen, J., Green-Church, K.B., Nichols, K.K., 2010. Shotgun lipidomic analysis of human meibomian gland secretions with electrospray ionization tandem mass spectrometry. Invest. Ophthalmol. Vis. Sci. 51, 6220–6231.

Chen, J., Green, K.B., Nichols, K.K., 2013. Quantitative profiling of major neutral lipid classes in human meibum by direct infusion electrospray ionization mass spectrometry. Invest. Ophthalmol. Vis. Sci. 54, 5730–5753.

Chen, J., Green, K.B., Nichols, K.K., 2016. Compositional analysis of wax esters in human meibomian gland secretions by direct infusion electrospray ionization mass spectrometry. Lipids 51, 1269–1287.

Chen, J., Nguyen, T.M.H., Zhi, W., 2025. Untargeted shotgun lipidomics via sequential precursor ion fragmentation on an Orbitrap mass spectrometer, 73rd ASMS Conference on Mass Spectrometry, Baltimore, MD, USA.

Chen, J., Nichols, K.K., 2018. Comprehensive shotgun lipidomics of human meibomian gland secretions using ms/msall with successive switching between acquisition polarity modes. J. Lipid Res. 59, 2223–2236.

Chen, J., Nichols, K.K., Wilson, L., Barnes, S., Nichols, J.J., 2019. Untargeted lipidomic analysis of human tears: A new approach for quantification of O-acyl-omega hydroxy fatty acids. The Ocular Surface 17, 347–355.

Chen, J., Panthi, S., 2019. Lipidomic analysis of meibomian gland secretions from the tree shrew: Identification of candidate tear lipids critical for reducing evaporation. Chem. Phys. Lipids 220, 36–48.

Davies, J.T., 1948. Monolayers of some diesters. Transactions of the Faraday Society 44, 909–913.

Davies, J.T., Rideal, E.K., 1963. Chapter 5-properties of monolayers, in: Davies, J.T., Rideal, E.K. (Eds.), Interfacial phenomena.

El-Hashani, A., Abuissa, H., Riheel, R., 2025. Benzalkonium chloride micellization: Salt and temperature dependence (conductivity, surface tension). Scientiae Radices 4, 181–201.

Ende, M., Spiteller, G., 1982. Contaminants in mass spectrometry. Mass Spectrom. Rev. 1, 29–62.

Flory, P.J., Vrij, A., 1963. Melting points of linear-chain homologs. The normal paraffin hydrocarbons. J. Am. Chem. Soc. 85, 3548–3553.

Flouda, P., Stryutsky, A.V., Buxton, M.L., Adstedt, K.M., Bukharina, D., Shevchenko, V.V., Tsukruk, V.V., 2022. Reconfiguration of Langmuir monolayers of thermo-responsive branched ionic polymers with LCST transition. Langmuir 38, 12070–12081.

Fodran, T., Fodran, P., Minnaard, A., Chlpík, J., Cirak, J., 2016. Thermomechanical properties of Langmuir-Blodgett monolayers of (R)-and (S)-tuberculostearic acid, The 22nd International Conference on Applied Physics of Condensed Matter, Štrbské Pleso, Slovak Republic.

Georgiev, G.A., Gh, M.S., Romano, J., Dias Teixeira, K.L., Struble, C., Ryan, D.S., Sia, R.K., Kitt, J.P., Harris, J.M., Hsu, K.-L., Libby, A., Odrich, M.G., Suárez, T., McKown, R.L., Laurie, G.W., 2021. Lacritin proteoforms prevent tear film collapse and maintain epithelial homeostasis. J. Biol. Chem. 296, 100070.

Georgiev, G.A., Kutsarova, E., Jordanova, A., Krastev, R., Lalchev, Z., 2010. Interactions of meibomian gland secretion with polar lipids in Langmuir monolayers. Colloids and Surfaces B: Biointerfaces 78, 317–327.

Georgiev, G.A., Yokoi, N., Koev, K., Kutsarova, E., Ivanova, S., Kyumurkov, A., Jordanova, A., Krastev, R., Lalchev, Z., 2011. Surface chemistry study of the interactions of benzalkonium chloride with films of meibum, corneal cells lipids, and whole tears. Invest. Ophthalmol. Vis. Sci. 52, 4645–4654.

Glasgow, B.J., Abduragimov, A.R., Farahbakhsh, Z.T., Faull, K.F., Hubbell, W.L., 1995. Tear lipocalins bind a broad array of lipid ligands. Curr. Eye Res. 14, 363–372.

Goldstein, M.H., Silva, F.Q., Blender, N., Tran, T., Vantipalli, S., 2022. Ocular benzalkonium chloride exposure: Problems and solutions. Eye 36, 361–368.

Guaus, E., Torrent-Burgués, J., 2018. The Langmuir technique applied to the study of natural tears. BioNanoScience 8, 559–565.

Han, X., Gross, R.W., 2005. Shotgun lipidomics: Electrospray ionization mass spectrometric analysis and quantitation of cellular lipidomes directly from crude extracts of biological samples. Mass Spectrom. Rev. 24, 367–412.

Howland, H.C., Merola, S., Basarab, J.R., 2004. The allometry and scaling of the size of vertebrate eyes. Vision Res. 44, 2043–2065.

Kahook, M.Y., Rapuano, C.J., Messmer, E.M., Radcliffe, N.M., Galor, A., Baudouin, C., 2024. Preservatives and ocular surface disease: A review. The Ocular Surface 34, 213–224.

Kato, Y., Yagi, H., Kaji, Y., Oshika, T., Goto, Y., 2013. Benzalkonium chloride accelerates the formation of the amyloid fibrils of corneal dystrophy-associated peptides. J. Biol. Chem. 288, 25109–25118.

Keller, B.O., Sui, J., Young, A.B., Whittal, R.M., 2008. Interferences and contaminants encountered in modern mass spectrometry. Anal. Chim. Acta 627, 71–81.

Khanom, M., Borchman, D., Jaeger, V.W., 2026. A detailed molecular model of the tear film lipid layer. The Journal of Physical Chemistry B 130, 7375–7387.

King-Smith, P.E., Bailey, M.D., Braun, R.J., 2013. Four characteristics and a model of an effective tear film lipid layer (TFLL). Ocul Surf 11, 236–245.

King-Smith, P.E., Nichols, J.J., Braun, R.J., Nichols, K.K., 2011. High resolution microscopy of the lipid layer of the tear film. Ocul Surf 9, 197–211.

Knothe, G., Dunn, R.O., 2009. A comprehensive evaluation of the melting points of fatty acids and esters determined by differential scanning calorimetry. Journal of the American Oil Chemists’ Society 86, 843–856.

Kolattukudy, P.E., Rogers, L.M., Nicolaides, N., 1985. Biosynthesis of lipids by bovine meibomian glands. Lipids 20, 468–474.

Kotz, J.C.T., Paul M.; Townsend, John R.; Treichel, David A., 2019. Intermolecular forces and liquids, in: Kotz, J.C.T., Paul M.; Townsend, John R.; Treichel, David A. (Ed.), Chemistry & chemical reactivity, 10th ed. Cengage Learning, Boston, MA, USA, pp. 490–518.

Kunnen, C.M., Brown, S.H., Lazon de la Jara, P., Holden, B.A., Blanksby, S.J., Mitchell, T.W., Papas, E.B., 2016. Influence of meibomian gland expression methods on human lipid analysis results. Ocul Surf 14, 49–55.

La Mer, V.K., Aylmore, L.A., 1962. Evaporation resistance as a sensitive measure of the purity and molecular structure of monolayers. Proc. Natl. Acad. Sci. U. S. A. 48, 316–324.

La Mer, V.K., Healy, T.W., 1965. Evaporation of water: Its retardation by monolayers: Spreading a monomolecular film on the surface is a tested and economical means of reducing water loss. Science 148, 36–42.

Lam, S.M., Tong, L., Reux, B., Lear, M.J., Wenk, M.R., Shui, G.H., 2013. Rapid and sensitive profiling of tear wax ester species using high performance liquid chromatography coupled with tandem mass spectrometry. J. Chromatogr. A 1308, 166–171.

Leiske, D., Leiske, C., Leiske, D., Toney, M., Senchyna, M., Ketelson, H., Meadows, D., Fuller, G.G., 2012a. Temperature-induced transitions in the structure and interfacial rheology of human meibum. Biophys. J. 102, 369–376.

Leiske, D.L., Miller, C.E., Rosenfeld, L., Cerretani, C., Ayzner, A., Lin, B., Meron, M., Senchyna, M., Ketelson, H.A., Meadows, D., Srinivasan, S., Jones, L., Radke, C.J., Toney, M.F., Fuller, G.G., 2012b. Molecular structure of interfacial human meibum films. Langmuir 28, 11858–11865.

Leiske, D.L., Raju, S.R., Ketelson, H.A., Millar, T.J., Fuller, G.G., 2010. The interfacial viscoelastic properties and structures of human and animal meibomian lipids. Exp. Eye Res. 90, 598–604.

Li, S., Du, L., Zhang, Q., Wang, W., 2018. Stabilizing mixed fatty acid and phthalate ester monolayer on artificial seawater. Environmental Pollution 242, 626–633.

Liu, J., Deng, W., Yu, M., Wen, R., Yao, S., Chen, B., 2017. Rapid analysis of benzalkonium chloride using paper spray mass spectrometry. J. Pharm. Biomed. Anal. 145, 151–157.

McCulley, J.P., Shine, W., 1997. A compositional based model for the tear film lipid layer. Trans. Am. Ophthalmol. Soc. 95, 79–88; discussion 88-93.

Millar, T.J., King-Smith, P.E., 2012. Analysis of comparison of human meibomian lipid films and mixtures with cholesteryl esters in vitro films using high resolution color microscopy. Invest. Ophthalmol. Vis. Sci. 53, 4710–4719.

Mudgil, P., Millar, T.J., 2008. Adsorption of apo- and holo-tear lipocalin to a bovine meibomian lipid film. Exp. Eye Res. 86, 622–628.

Mudgil, P., Millar, T.J., 2011. Surfactant properties of human meibomian lipids. Invest. Ophthalmol. Vis. Sci. 52, 1661–1670.

Nakahara, H., Nakamura, S., Nakamura, K., Inagaki, M., Aso, M., Higuchi, R., Shibata, O., 2005. Cerebroside Langmuir monolayers originated from the echinoderms: II. Binary systems of cerebrosides and steroids. Colloids and Surfaces B: Biointerfaces 42, 175–185.

Nicolaides, N., Kaitaranta, J.K., Rawdah, T.N., Macy, J.I., Boswell, F.M., 3rd, Smith, R.E., 1981. Meibomian gland studies: Comparison of steer and human lipids. Invest. Ophthalmol. Vis. Sci. 20, 522–536.

Oliveira, O.N., Jr., Caseli, L., Ariga, K., 2022. The past and the future of Langmuir and Langmuir–Blodgett films. Chem. Rev. 122, 6459–6513.

Petrov, J.G., Polymeropoulos, E.E., Möhwald, H., 2000. Langmuir monolayers with fluorinated groups in the hydrophilic head. 1. Comparison of trifluoroethyl behenate and ethyl behenate monolayers: molecular models, mechanical properties, stability. Langmuir 16, 7411–7420.

Petrov, P.G., Thompson, J.M., Rahman, I.B.A., Ellis, R.E., Green, E.M., Miano, F., Winlove, C.P., 2007. Two-dimensional order in mammalian pre-ocular tear film. Exp. Eye Res. 84, 1140–1146.

Politano, G.G., Versace, C., 2023. Spectroscopic ellipsometry: Advancements, applications and future prospects in optical characterization. Spectroscopy Journal 1, 163–181.

Riedlová, K., Saija, M.C., Olżyńska, A., Jurkiewicz, P., Daull, P., Garrigue, J.-S., Cwiklik, L., 2023. Influence of baks on tear film lipid layer: In vitro and in silico models. Eur. J. Pharm. Biopharm. 186, 65–73.

Rosenfeld, L., Cerretani, C., Leiske, D.L., Toney, M.F., Radke, C.J., Fuller, G.G., 2013. Structural and rheological properties of meibomian lipid. Invest. Ophthalmol. Vis. Sci. 54, 2720–2732.

Sadgrove, N.J., Jones, G.L., 2019. From petri dish to patient: Bioavailability estimation and mechanism of action for antimicrobial and immunomodulatory natural products. Front. Microbiol. Volume 10 - 2019.

Saville, J.T., Zhao, Z., Willcox, M.D., Ariyavidana, M.A., Blanksby, S.J., Mitchell, T.W., 2011. Identification of phospholipids in human meibum by nano-electrospray ionisation tandem mass spectrometry. Exp. Eye Res. 92, 238–240.

Schuett, B.S., Millar, T.J., 2012. Lipid component contributions to the surface activity of meibomian lipids. Invest. Ophthalmol. Vis. Sci. 53, 7208–7219.

Smaby, J.M., Kulkarni, V.S., Momsen, M., Brown, R.E., 1996. The interfacial elastic packing interactions of galactosylceramides, sphingomyelins, and phosphatidylcholines. Biophys. J. 70, 868–877.

Smith, E.C., Crane, J.M., Laderas, T.G., Hall, S.B., 2003. Metastability of a supercompressed fluid monolayer. Biophys. J. 85, 3048–3057.

Tiffany, J.M., Winter, N., Bliss, G., 1989. Tear film stability and tear surface tension. Curr. Eye Res. 8, 507–515.

Trevorah, R.M., Stubb, H., Viljanen, M., Mäkinen, H., Viitaja, T., Sevón, J., Konovalov, O., Jankowski, M., Hemmerle, A., Ekholm, F.S., Svedström, K.J., 2025. One step closer to a molecular level understanding of the tear film lipid layer by surface x-ray scattering. Langmuir 41, 23353–23361.

Trevorah, R.M., Viljanen, M., Viitaja, T., Stubb, H., Sevón, J., Konovalov, O., Jankowski, M., Fontaine, P., Hemmerle, A., Raitanen, J.-E., Ekholm, F.S., Svedström, K.J., 2024. New insights into the molecular structure of tear film lipids revealed by surface x-ray scattering. The Journal of Physical Chemistry Letters 15, 316–322.

Tsukruk, V.V., Singamaneni, S., 2012. Scanning probe microscopy of soft matter: Fundamentals and practices. Wiley-VCH, Weinheim.

Vargaftik, N.B., Volkov, B.N., Voljak, L.D., 1983. International tables of the surface tension of water. Journal of Physical and Chemical Reference Data 12, 817–820.

Willcox, M.D.P., Argüeso, P., Georgiev, G.A., Holopainen, J.M., Laurie, G.W., Millar, T.J., Papas, E.B., Rolland, J.P., Schmidt, T.A., Stahl, U., Suarez, T., Subbaraman, L.N., Uçakhan, O.Ö., Jones, L., 2017. TFOS DEWS II Tear Film Report. The Ocular Surface 15, 366–403.

Wnętrzak, A., Lipiec, E., Łątka, K., Kwiatek, W., Dynarowicz-Łątka, P., 2014. Affinity of alkylphosphocholines to biological membrane of prostate cancer: Studies in natural and model systems. The Journal of Membrane Biology 247, 581–589.

Yoshida, M., Yamaguchi, M., Sato, A., Tabuchi, N., Kon, R., Iimura, K.-i., 2019. Role of endogenous ingredients in meibum and film structures on stability of the tear film lipid layer against lateral compression. Langmuir 35, 8445–8451.

Züllig, T., Trötzmüller, M., Köfeler, H.C., 2020. Lipidomics from sample preparation to data analysis: A primer. Anal. Bioanal. Chem. 412, 2191–2209.

