## Supplementary figures and images for "Surface Chemistry and Interfacial Biophysics of Meibum and Meibum—Benzalkonium Chloride Films: Implications for Tear Film Lipid Layer"

### Suppl Fig. 1

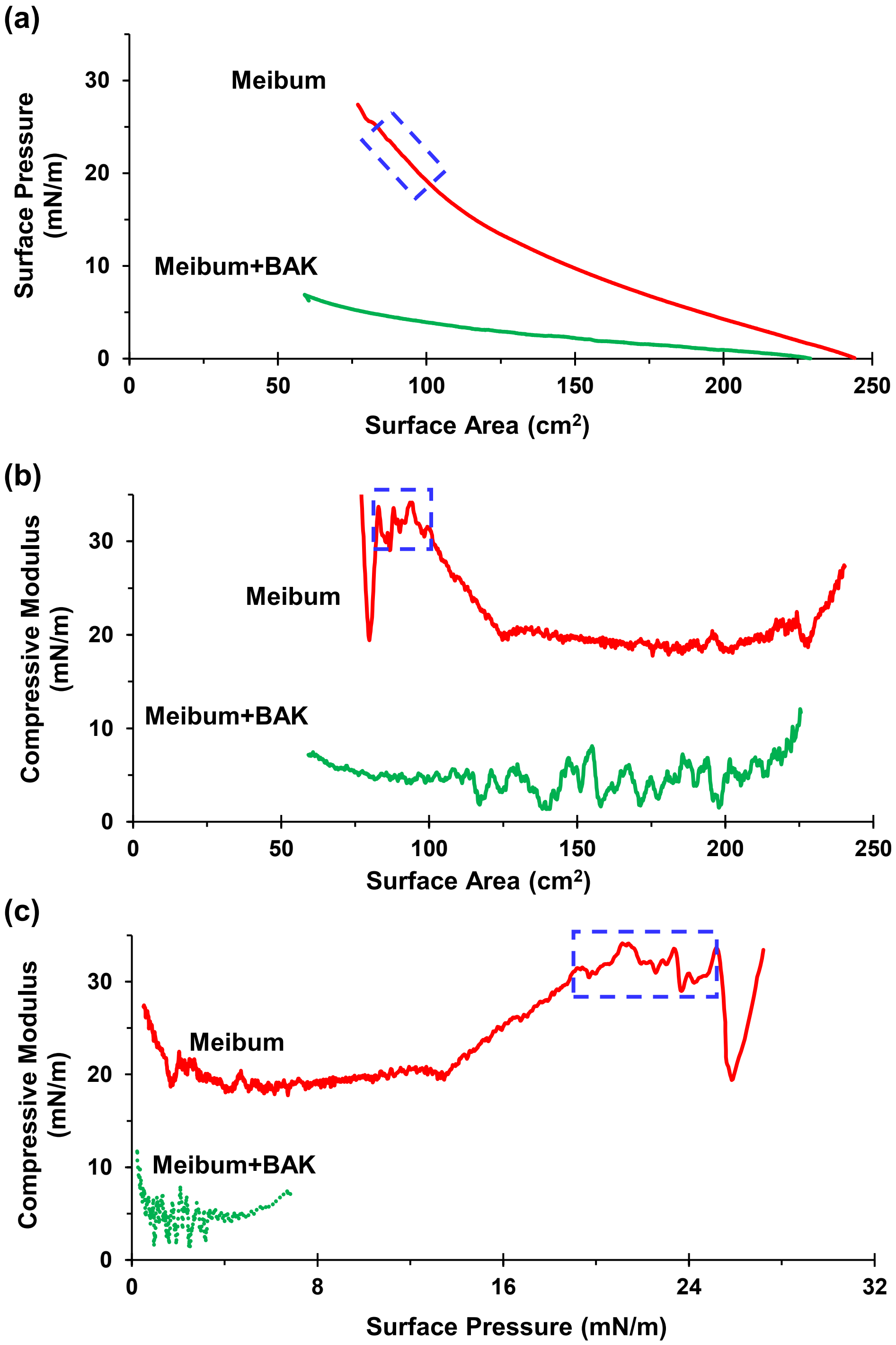
